# The *ecnA* antitoxin is not only important for human pathogens: Evidences of its role in the plant pathogen *Xathomonas citri* subsp. *citri*

**DOI:** 10.1101/505081

**Authors:** Laís Moreira Granato, Simone Cristina Picchi, Maxuel de Oliveira Andrade, Paula Maria Moreira Martins, Marco Aurélio Takita, Marcos Antonio Machado, Alessandra Alves de Souza

**Author notes:** Address correspondence to Dra. Alessandra Alves de Souza Mailing address: Centro de Citricultura Sylvio Moreira/IAC, Rodovia Anhanguera Km 158, Cordeirópolis SP, Brazil, 13490-970.

## Abstract

*Xanthomonas citri* subsp. *citri* causes citrus canker disease worldwide in most commercial varieties of citrus. Its transmission occurs mainly through wind-driven rain. Once on the leaf *X. citri* can epiphytically survive forming biofilm, which enhances persistence of the bacteria to different environmental stresses and play an important role in the early stages of host infection. Therefore, the study of genes involved in biofilm formation has been an important step towards the understanding of the bacterial strategy to survive and infect the plant host. In this work we show that e*cnAB* a Toxin-Antitoxin (TA) system, previously identified only in human bacterial pathogen, is conserved in many *Xanthomonas* spp. In general TA systems consist of a pair of genes in operon that encodes a stable toxin and an unstable antitoxin that, under normal conditions, binds to the toxin and blocks its activity. On the other hand, under stress the antitoxin is degraded, allowing the toxin to act decreasing cell growth and metabolism. When normal growth conditions are re-established, the antitoxin is produced, blocking the toxin and allowing the cells to grow. Thus, this mechanism represents an important bacterial strategy of survival under stress conditions. In this work, we show that in *X. citri ecnAB* is regulated by quorum sensing and it is involved in important processes such as biofilm formation, EPS production, and motility. In addition, we show that *ecnAB* plays a role in *X. citri* survival and virulence in plant host.

**IMPORTANCE:** Very little is known about TA systems in phytopathogenic bacteria. The *ecnAB* in special has been only studied in bacterial human pathogens. Here we showed that it is present in a wide range of the phytopathogen *Xanthomonas spp*., and moreover this is the first work that investigated the functional role of this TA system in *Xanthomonas citri* biology suggesting an important new role in adaptation and survival with implications in the bacterial pathogenicity.

## INTRODUCTION

Phytopathogenic bacteria such as *Xanthomonas spp.* cause huge damages in many different crops such as brassicas, rice, cassava, tomato and citrus (1). *Xanthomonas citri* subsp. *citri* (*X. citri*) causes citrus canker disease worldwide in most commercial varieties of citrus (2), for this reason there was an increase in the number of works focusing on the identification of genes involved with the mechanism of pathogenicity and plant-pathogen interaction aiming the development of strategies to control *X. citri* (3–8) *X. citri* is disperse by wind-driven rain and then it adheres to the plant leaf surfaces (9). The stable adhesion to the host is the first step towards biofilm formation (10), which enhances the epiphytic persistence of the bacteria on host leaves, plays an important role in the early stages of infection (11), and confers resistance to different environmental stresses, including UV radiation, salt, osmotic challenge, desiccation and H_2_O_2_ (12–14). Mutants of genes involved with biofilm formation are less virulent (15–18), demonstrating that *X. citri* biofilm is also associated with virulence. Thus, the study of genes involved in biofilm formation has been an important step to understand the bacterial strategy to survive and infect the plant host.

One of the genetic mechanism involved in biofilm formation is the quorum sensing, which is a refined process of cellular communication (19). Mutations in genes involved with quorum-sensing like *rpfF, rpfG*, and *rpfC* in *Xanthomonas* led to the reduction of the synthesis of virulence factors, reduction of xanthan gum production and biofilm development (18, 20, 21). Thus, the genes regulated by quorum sensing are important to be understood since they are directly involved in features that allow the success of bacterial survival and virulence in the host (18). In fact, the transcriptome analysis of the *rpfF* mutant revealed a set of genes involved in chemotaxis and motility, adhesion, stress tolerance, regulation, transport, and detoxification (18). Many of these genes are extensively studied (18, 22, 23), however there are genes whose functions in *X. citri* are still unknown such as the e*cnAB* toxin/antitoxin (TA) system (XAC_RS20190/XAC_RS20185) despite its important role in *E. coli* has already been demonstrated (24, 25). The *E. coli ecnAB* locus encodes the synthesis of two small lipoproteins that are part the entericidin locus TA system, because of hypothesized similarities to a plasmid addiction module, *mazEF*, that has been studied in *E. coli* (24). EcnA was designated as the antitoxin that counteracts the EcnB toxin, whose overproduction results in a more pronounced bacterial lysis than that observed in the co-expression of both *ecnA* and *ecnB* (25). Different functions have been proposed for the *ecnAB* TA system, including programmed cell death (24), adherence in the respiratory tract of humans (26), and its requirement for the probiotic functions of *Enterobacter* sp. strain C6-6 (25).

Most of the studies regarding TA systems are related to human-associated bacteria, and only a few have addressed TAs from phytopathogens (27). Indeed TA systems have been implicated in induction of persister cells and consequently the maintenance of phytopathogen virulence (28). In Xanthomonas, just one work showed the quantitative and qualitative differences among the type II TA systems (29) however no gene had the function characterized. In this work, the importance of the *ecnAB* locus is demonstrated in *Xanthomonas* for the first time and we suggest that this locus is involved in the regulation of genes related with biofilm formation, motility and cell survival, thereby affecting its virulence in the plant host.

## MATERIALS AND METHODS

### Bacterial strains, growth conditions and plasmids

The bacterial strains and plasmids employed in this study are described in Table 1. *E. coli* DH5α cells were routinely grown at 37 °C in Luria-Bertani (LB) medium (1% (w/v) tryptone, 0.5% (w/v) yeast extract and 1 % (w/v) sodium chloride, pH 7.5) shaking at 180 rpm or on plates. *Xanthomonas citri* subsp. *citri* strain 306 (30) and derivatives were grown at 28 °C in NBY nutrient medium (0.5 % (w/v) peptone, 0.3 % (w/v) meat extract, 0.2 % (w/v) yeast extract, 0.2 % (w/v) K_2_HPO_4_, 0.05% (w/v) KH_2_PO_4_, pH 7.2) shaking at 180 rpm or on 1.2 % (w/v) agar solid media. Antibiotics were used at the following concentrations: ampicillin (Ap) 100 μg/mL, kanamycin (Km) 50 μg/mL, gentamycin (Gm) 5 μg/mL.

**Table 1:** Strains and plasmids used in this study

| Strains | Characteristics | Reference or source |
| --- | --- | --- |
| *Escherichia coli* Dh5α | F<sup>-</sup> ϕ80dlacZ hM15 h (*lacZYA*-argF) U169<br>*endA1 deoR recA1 hsdR17*(r<sub>K</sub><sup>-</sup> m<sub>K</sub><sup>+</sup>) *phoA*<br>*supE44* λ<sup>-</sup> *thi-1* *gyrA96* | (31) |
| *Xanthomonas citri* subsp. *citri*<br>306<br>(*X. citri*) | Syn. *X. axonopodis* pv. *citri* strain 306; wild-type, Ap<sup>r</sup> | (30, 32) |
| B3/*ecnA*::Tn5 (*ecnA*<sup>-</sup>) | *ecnA*<sup>-</sup> (XAC_RS20190), Km<sup>r</sup> | (33) |
| *ecnA*::Tn5_c (*ecnA*<sup>+</sup>) | *ecnA*<sup>+</sup> (contained in pUFR053, Km<sup>r</sup> Gm<sup>r</sup>) | This study |
| Δ*rpfF* (*rpfF*<sup>-</sup>) | Δ*rpfF* (XAC_RS09555), Ap<sup>r</sup> | This study |
| Plasmids |  |  |
| pNPTS138 | Suicide vector for generation of gene<br>knockouts, *sacB* and Km<sup>r</sup> | (34) |
| pNPTS-Δ*rpfF* | pNPTS138 derivative for generation of *flaA*<br>knockout, *sacB* and Km<sup>r</sup> | This study |
| pUFR053 | IncW Mob<sup>+</sup> mob (P) *lacZ*α<sup>+</sup> Par<sup>+</sup>, Cm<sup>r</sup>, Gm<sup>r</sup>,<br>Km<sup>r</sup>, shuttle vector | (35) |
| p53_*ecnA* | *ecnA* gene cloning on pUFR053 | This study |

Ap<sup>r</sup>, Km<sup>r</sup>, Gm<sup>r</sup> indicate resistance to ampicillin, kanamycin, gentamicin, respectively

### Molecular Biology

Bacterial genomic and plasmid DNA were isolated using a Wizard genomic DNA purification kit and a Wizard miniprep DNA purification system according to the manufacturer’s instructions (Promega, Madison, WI, USA). Concentration and purity of the DNA were determined by using a Nanodrop ND-1000 spectrophotometer (NanoDrop Technologies, Wilmington, DE, USA). PCR and DNA electrophoresis was performed as described (36) with *Pfu* DNA polymerase (Promega Corporation, Madison, WI). Restriction digestions and DNA ligations were performed according to the manufacturer’s instructions (New England Biolabs, USA).

### In-frame deletion of *rpfF*

To construct the *rpfF* deletion mutant, approximately 1 kb of the upstream and downstream regions of the *rpfF* gene (XAC_RS09555) were amplified by PCR from *X. citri* genomic DNA. The first fragment was amplified with the oligos sequences Fwd1-5’-TGAGCAAGCTTCCGCGCGACATGCCAGGTTTCG-3’ and Rv1-5’-TCGTGCT CCATGGGTTTGATCCTGGTAAGGCCGCG-3’ and the second fragment by using the oligos sequences Fwd2-5’-TGCGCGTCGGCCATGGGCGTCGCTGATTTTTGAT-3’ and Rv2-5’-TGCAGGATGCCGACACCACCGCCAGATCTCCGG-3’. Fragments were ligated to produce an in-frame deletion. This sequence was then cloned into the *HindIII* and *BglII* restriction sites of the pNPTS138 suicide vector, thus generating the plasmid pNPTS-rpfF. This plasmid was introduced into *X. citri* cells by electroporation, and the wild-type copy was replaced by the deleted version after two recombination events as described previously (34).

### Quorum sensing regulation

To verify if the DSF-mediated quorum sensing system is involved in the regulation of *ecnAB*, a quantitative reverse transcription PCR assay was performed using the *rpfF* deletion mutant. The RNA extraction from the wild-type and *rpfF* mutant strains was done using RNeasy Plant minikit (Qiagen) and RNA quality and concentration was assessed by gel electrophoresis and Nanodrop (Thermo Scientific). cDNA was synthesized with GoScript Reverse Transcription System (Promega) and qPCR was realized using GoTaq SYBR green (Promega) and followed the manufacturer instructions in a 7500 Fast Real-Time PCR system (Applied) and the output data were analyzed by fold change calculation (2^-ΔΔCT^). Gene expression was estimated using primers for the genes from the *ecnAB* locus. *gumB* was used as control, since it is negatively regulated by *rpfF* and 16S rRNA was also used as the endogenous control (Table 2). Three independent experiments were performed and statistical significance among the means was calculated using ANOVA followed by Tukey test (*P* < 0.05).

**Table 2:**
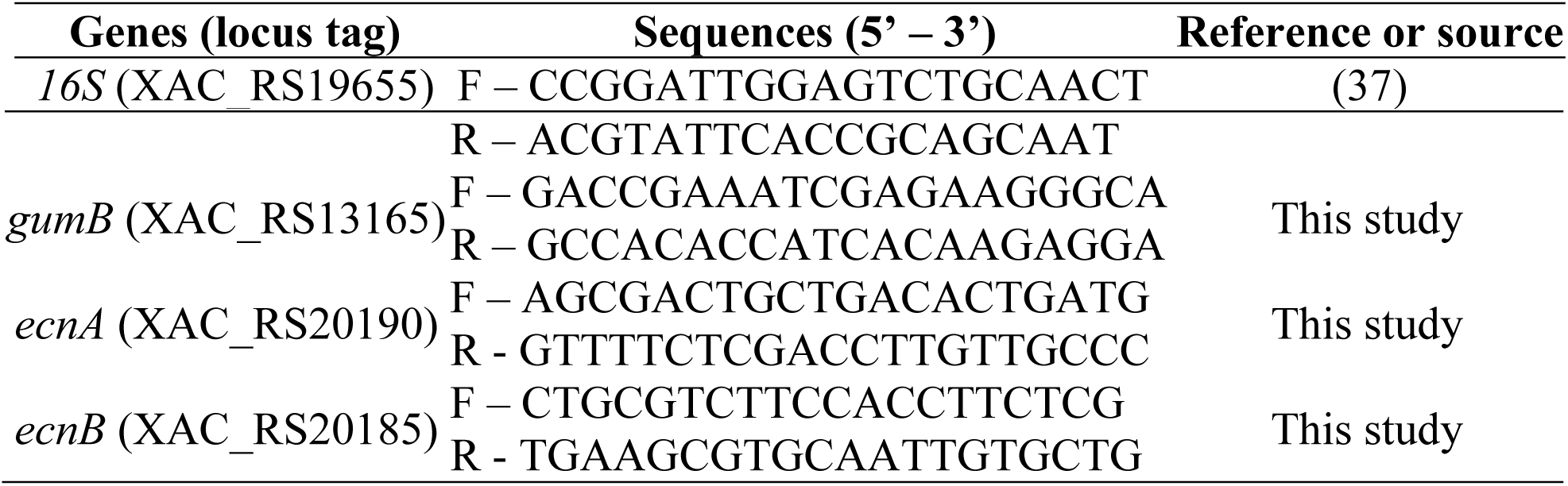
Primers used for qRT-PCR

### Construction of the complementation vector p53_*ecnA*

To complement the *ecnA*::Tn5 mutant, a 153 bp DNA fragment harboring the *ecnA* ORF was amplified by PCR using total DNA obtained from the *X. citri* wild-type strain 306 as the template and the specific primer pair *ecnA*_p53_Fwd (5′ TCGAAGCTTCAGCGAATGCAGGTGTTC 3′) and *ecnA*_p53_Rv (5′ GCGATGTGACGGATCAGCGAATTCCTG 3′). The PCR-amplified fragment was digested with *HindIII* and *EcoRI* at the recognition sites designed in the primer sequences and this processed fragment was ligated into pUFR053 (35), resulting in p53_*ecnA* (Table 1). The construction was confirmed using sequencing and the plasmid was transferred into *ecnA::Tn5* using electroporation. The cells were selected on NBY solid media using gentamicin, resulting in the strain *ecnA*::Tn5_c (*ecnA*+).

### Biofilm formation assays

Biofilms formation on polystyrene (Nunc™ 96 well-plates) and glass surfaces was examined as described previously (38) with modifications. The wild-type, mutant and complemented cells were grown into NBY overnight at 28 °C with shaking (180 rpm). The optical density (OD) at 600 nm was adjusted to 0.1 (UV/Vis Spectrometer Lambda Bio; Perkin Elmer) using fresh NBY plus 1.0 % (w/v) D-glucose. A 200 µL portion of adjusted bacteria was placed in each well of the 96-well plates and 1.5mL portion of adjusted bacteria was placed in the glass tubes and both were incubated at 28 °C without agitation. After 48 h, bacterial growth was checked, and the culture was removed using a pipette. After repetitive water washes, the adhered cells were stained with 0.1% (w/v) CV for 30 min at room temperature. The unbound stain was removed by water washing, and the CV-stained cells were solubilized in ethanol. The samples on plates were measured at 590 nm using microplate reader 3550 (Bio-Rad) and normalized with bacterial growth (OD=600nm). Both tests were repeated three times independently with six replicates each. The means were statistically analysed using one-way analysis of variance (ANOVA) and Tukey test (*P* < 0.05).

Assays of biofilm formation on leaf surfaces were performed as previously described (39), with modifications. Briefly, 20 μL volume of each bacterial suspension (10^4^ CFU/mL) was dropped onto the leaves abaxial surfaces of sweet orange leaves. The leaves were maintained at 28 °C in a humidified chamber. The biofilm formation was visualized on the leaf surfaces using crystal violet staining. Leaf discs from staining spots were excised, dissolved in 1 mL ethanol:acetone (70:30, v/v) and quantified by measuring the optical density at 590 nm. Data from both experiments were statistically analysed using ANOVA and Tukey test (*P* < 0.05), and values are expressed as the means ± standard deviations.

### EPS quantification

The bacteria strains were cultivated overnight in NBY medium at 28°C with shaking at 200 rpm. The optical density (OD) at 600 nm was adjusted to 0.05 (UV/Vis Spectrometer Lambda Bio; Perkin Elmer) using fresh NBY plus 1.0 % (w/v) D-glucose. After 72 h the cells were pelleted by centrifugation (4456 g for 6 min). The supernatant was transferred to another tube, and 5 volumes of 100% ethanol were added. The crude xanthan was collected using a glass rod and placed on a Petri dish to dry at 60°C for 48 h. The dried xanthan was weighed, and the values were expressed as the means ± standard deviations. Data were statistically analysed using ANOVA and Tukey test (*P* < 0.05).

### Motility assays

Motility assays were performed as previously described (40). Bacteria were grown overnight in NBY medium, and 3 µL of bacterial culture (OD600 0.3) was then spotted onto a plate containing SB medium plus 0.5 % (wt/v) agar (Difco, Franklin Lakes, NJ) (41) for the sliding motility tests or NYGB medium 0.25 % (wt/v) agar (42) for the swimming motility tests. Plates were incubated at 28 °C for 48 h. The diameters of the circular halos that were occupied by the strains were measured, and the resulting values were taken to indicate the motility of *X. citri* strains. The experiments were repeated three times with three replicates each time. The diameter measurements were statistically analysed using ANOVA and Tukey test (*P* < 0.05), and the values are expressed as the means ± standard deviations of three independent experiments.

### Pathogenicity assay

*X. citri* 306 wild-type and *ecnA* mutant were grown in selective antibiotic NBY medium overnight at 28 °C, centrifuged at 4456 g and then resuspended in 10 mM potassium phosphate buffer (pH 7.0). Immature leaves from sweet orange (*Citrus sinensis* cv. ‘Bahia’) plants were inoculated with the bacterial suspensions by spraying or injected into the intercellular spaces of leaves with a needleless syringe (10^8^ CFU/mL). Phosphate buffer was used as the control in non-infected plants. Five plants were inoculated for each bacterial strain in each inoculation method. The plants were kept in the greenhouse at “Fazenda Santa Elisa” from Instituto Agronômico de Campinas/IAC (Campinas, SP), at a temperature of 28 ± 4 °C in high humidity for 30 days. At 7, 14, 21 and 30 days post-inoculation, three leaves with similar sizes were randomly collected from three different plants. To evaluate the spray assay, the leaves were immersed in 10 mL of phosphate buffer in Falcon flasks (50 mL). Bacterial cells were collected and then vortexed for three minutes to homogenize the tissue. In the infiltration assay, leaf discs (0.8 cm in diameter), randomly selected from the inoculated leaves, were ground in 1 mL of phosphate buffer. Bacterial numbers were determined in serial dilutions of the suspensions and then plated on NBY medium with the appropriate antibiotics. Colonies were counted after 48 hours of incubation at 28 °C. Disease symptoms severity was evaluated at 7, 14, 21 and 30 days after inoculation by three independents evaluators following the diagrammatic scale for citrus canker disease (43). Two independent experiments were performed. Statistical significance among the means was calculated with ANOVA and Tukey test (*P* < 0.05).

### Induction of oxidative stress

The cells were grown overnight in NBY medium at 28°C with shaking at 180rpm. The optical density (OD) at 600 nm was adjusted to 0.05 (UV/Vis Spectrometer Lambda Bio; Perkin Elmer) using fresh NBY. At this point, 1 mL aliquot was added into the 1.5 mL plastic tubes to which were supplemented with hydrogen peroxide to reach a 50 mM concentration. This concentration was chosen to induce oxidative stress in *E. coli* (44). These cells were kept at this stress for 10 min at approximately 28°C without agitation. The RNA extraction, RNA quality, cDNA synthesis and qPCR were performed as described above. Relative expression was evaluated using the 2^−ΔΔCT^ method using the 16S gene expression as endogenous control (table 2). The experiment was performed using three biological replicates and statistical significance among the means was calculated using ANOVA followed by Tukey test (*P* < 0.05).

## RESULTS

### *ecnA* encodes a probable antitoxin

The *ecnA* mutant (B3) was isolated from an EZ-Tn5 transposon mutagenesis library of *X. citri* strain 306 (33), which was identified among others with altered pattern in forming biofilm. Sequencing indicated that EZ-Tn5 was inserted between nucleotides 4 and 105 of XAC_RS20190 locus in the B3 mutant (Fig. 1A). The mutant was named *ecnA*::Tn5, since the transposon had the insertion located in the *ecnA* gene, which encodes a protein with similar identification to the EcnA protein, belonging to the entericidin locus in *E. coli*. To confirm the insertion, PCR using *ecnA* specific primers ecnA_F (5’ CAGCATTCCCACCGACAACATC 3’) and ecnA_R (5’ GCATGGGGTCGTTGGATATCGT 3’) was carried and resulted in a 0.52 kb amplicon when *X. citri* subsp. *citri* 306 genomic DNA was used as template and approximately 2.5 kb when the B3 mutant was used as template, which corresponds to the genomic fragment 1.938 kb plus the EZ-Tn5 transposon insertion (Fig. 1B).

**Figure 1.**
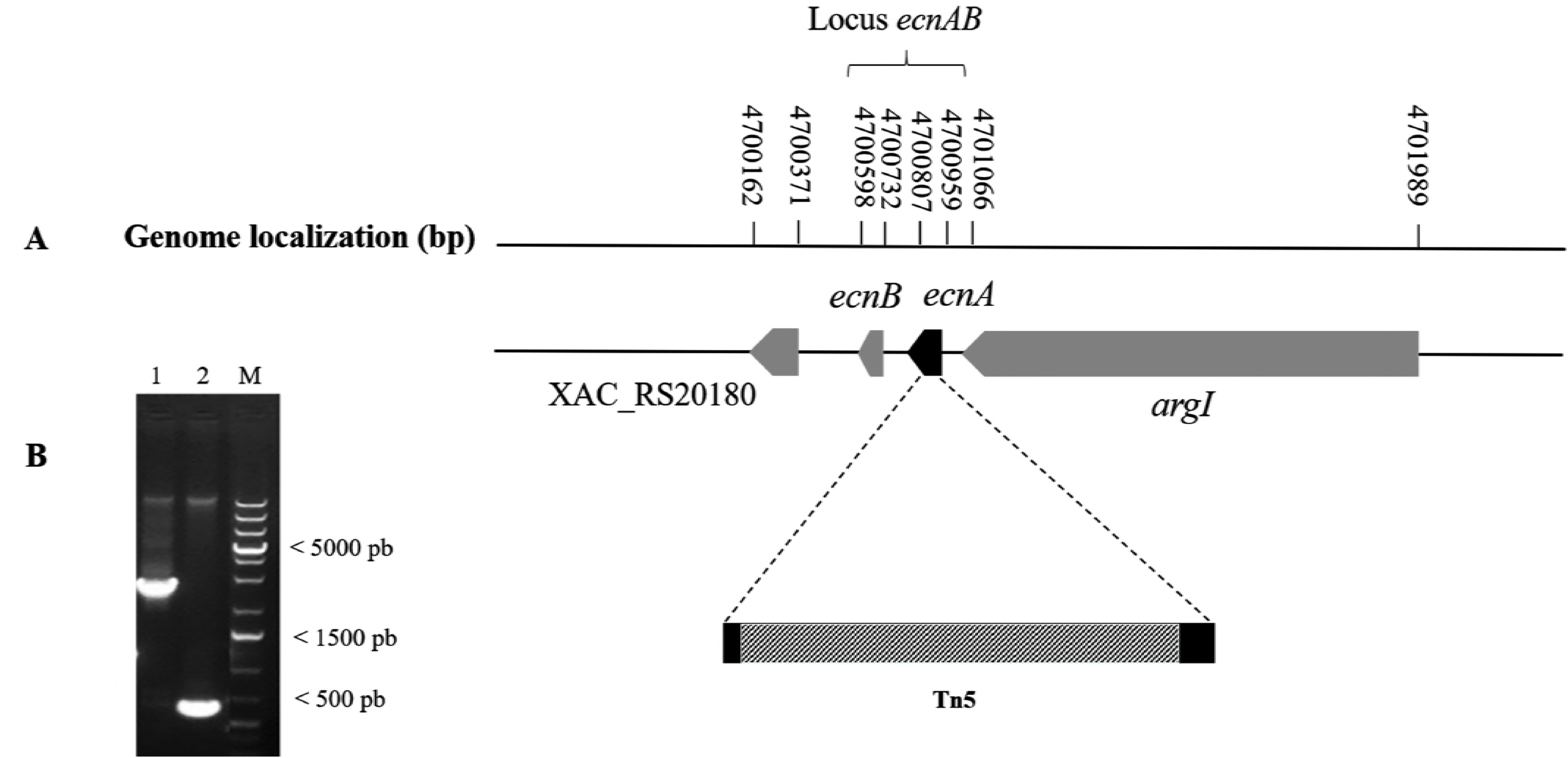
Sequence analysis of EZ-Tn5 insertion in the *ecnA* mutant strain. (A) Genomic location of *ecnA* on the *X. citri* chromosome showing the transposon insertion site between nucleotides 4 and 105 of *ecnA*. (B) PCR analysis confirming insertion of EZ-Tn5 in the *ecnA* gene: agarose gel electrophoresis of DNA amplified using primers ecnA_F and ecnA_R targeting a 500 pb region surrounding *ecnA* from the wild-type *X. citri* and B3 strains. Lane 1 - *X. citri*, lane 2 - B3 (*ecnA*::Tn5), M - Invitrogen 1 Kb Plus DNA size marker.

We verified that the *ecnAB* locus is conserved in other *Xanthomonas* species, including *X. fuscans* subsp. *fuscans, X. campestris* pv. vesicatoria, *X. campestris* pv. campestris, *X. oryzae* pv. oryzae, *X. oryzae* pv orizycola (Fig. 2).

**Figure 2.**
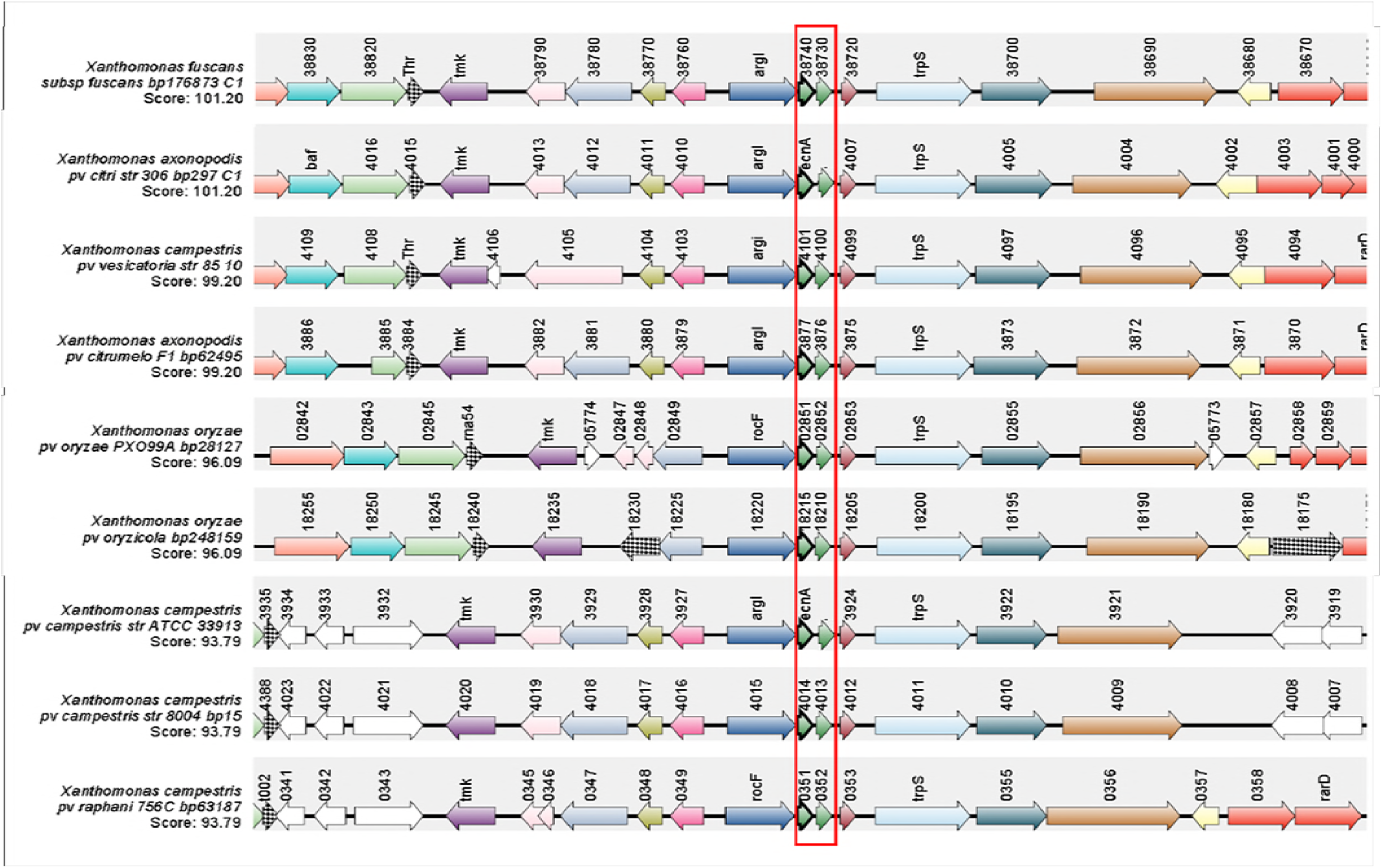
Synteny of the *ecnAB* locus. Genomic regions of different *Xanthomonas* spp. were aligned by the *ecnAB* locus, which is highlighted in red. The synteny shows that the *ecnAB* locus is conserved among *Xanthomonas* species.

### The *ecnAB* locus is regulated by *X. citri* quorum sensing and affects biofilm formation

It is known that *ecnA* transcription is regulated by quorum sensing (18). To confirm the effect of the quorum sensing regulatory components on expression of *X. citri ecnAB* system, we evaluated its expression in the *rpfF* mutant background, in which no DSF is produced (18). *gumB* was used as control, since it is known to be positively regulated by RpfF (45). We verified that both *ecnA* and *ecnB* were down-regulated in the *rpfF* mutant (Fig. 3). These results confirm that RpfF positively regulates *ecnAB* system. To assess the role of EcnA on biofilm we measured its development on abiotic and biotic surfaces. Intriguingly, contrary to *rpfF* mutant, significantly less biofilm formation was observed for the *ecnA* mutant in abiotic and biotic surfaces (Fig. 4). Taken together the results show that even though *ecnAB* are regulated by *rpfF*, the *ecnA* and *rpfF* mutants have a different behaviour regarding biofilm formation indicating that other components may play a role in this phenotype.

**Figure 3.**
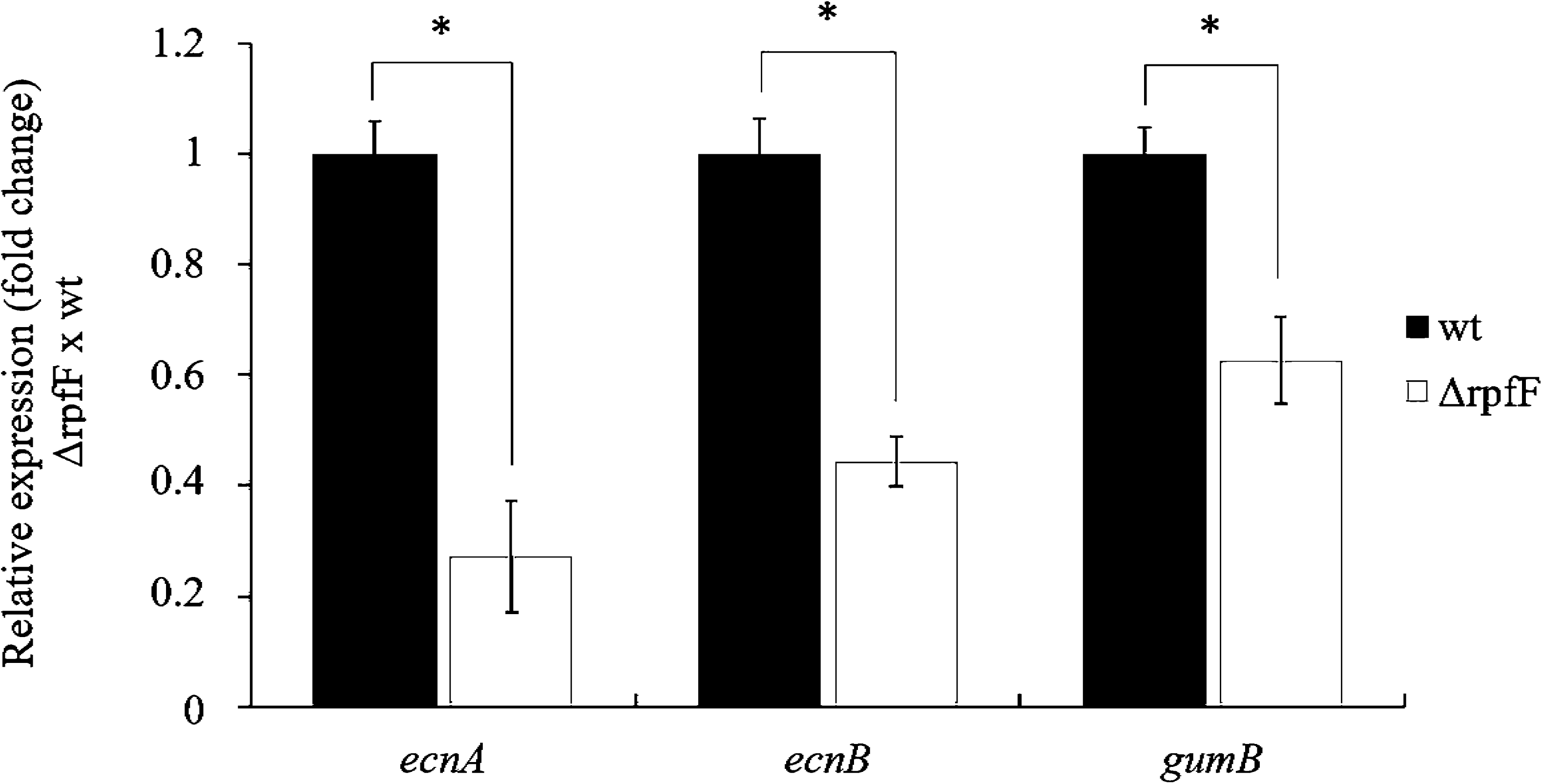
Gene expression analysis of *ecnAB* in *rpfF* mutant strain. Relative expression of the *ecnA, ecnB* and *gumB* genes in *rpfF* mutant compared to wild-type strain. The *gumB* was used as control to be positively regulated by RpfF (42). The transcript abundance was determined by real time RT-qPCR. Data are shown as the mean of three independent biological replicates, and error bars indicate the standard deviation of the mean. 16S rRNA was used as endogenous control. *indicates significant difference (*P* < 0.05) and ns indicates no significant.

**Figure 4.**
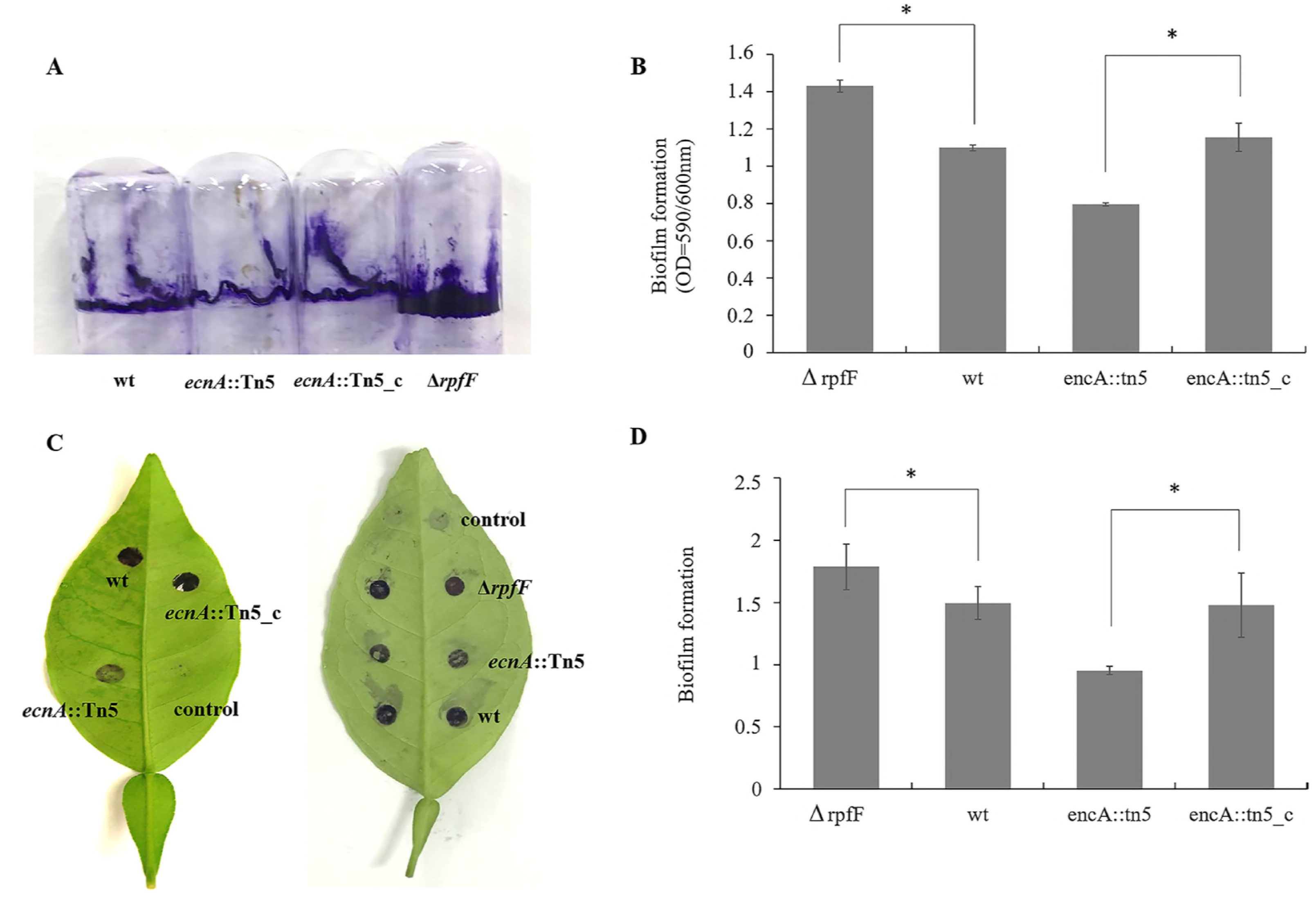
Biofilm formation by *X. citri ecnA* mutant and complemented strains. Biofilms formed on (A) abiotic and (C) biotic surfaces were stained with crystal violet and respectively quantified at OD=590nm (B and D). The values in B were normalized with bacterial growth (OD=600nm). Image of leaves is representative of six independent replicates. Error bars indicate the standard deviation of two independent experiments mean with six replicates each. *Indicates significant difference (*P* < 0.05) and ns indicates no significance. wt = wild-type *X. citri*, control = NBY medium plus 1.0 % (w/v) D-glucose without inoculation of *X. citri, ecnA*::Tn5 = *ecnA* mutant, *ecnA*::Tn5_c = complemented *ecnA* mutant.

### *ecnA* is involved in *X. citri* polysaccharide production and sliding motility

Because of the significant effect of the *ecnA* on biofilm formation, we evaluated if this feature was an effect of the alteration in exopolysaccharide production. Indeed, a significant reduction in EPS was observed in *ecnA* mutant (Fig. 5A) with a lower amount and dryer crude xanthan gum (Fig. 5B). These results suggest that the less biofilm formation observed in *ecnA* mutant could be related to the altered production of EPS. Besides its participation in biofilm, EPS has also a role in movement through sliding motility (11). Indeed, *ecnA* mutant show significant reduced diameters of the sliding motility compared to the wild-type and *ecnA*+ strains (Fig. 6). Swimming analyses were also performed, but no difference was observed between the mutant and wild-type strains (Fig. S1). Growth curve assays were performed, and no difference was observed between the strains (Fig. S2), indicating that the difference observed in the motility assays was not related to growth.

**Figure 5.**
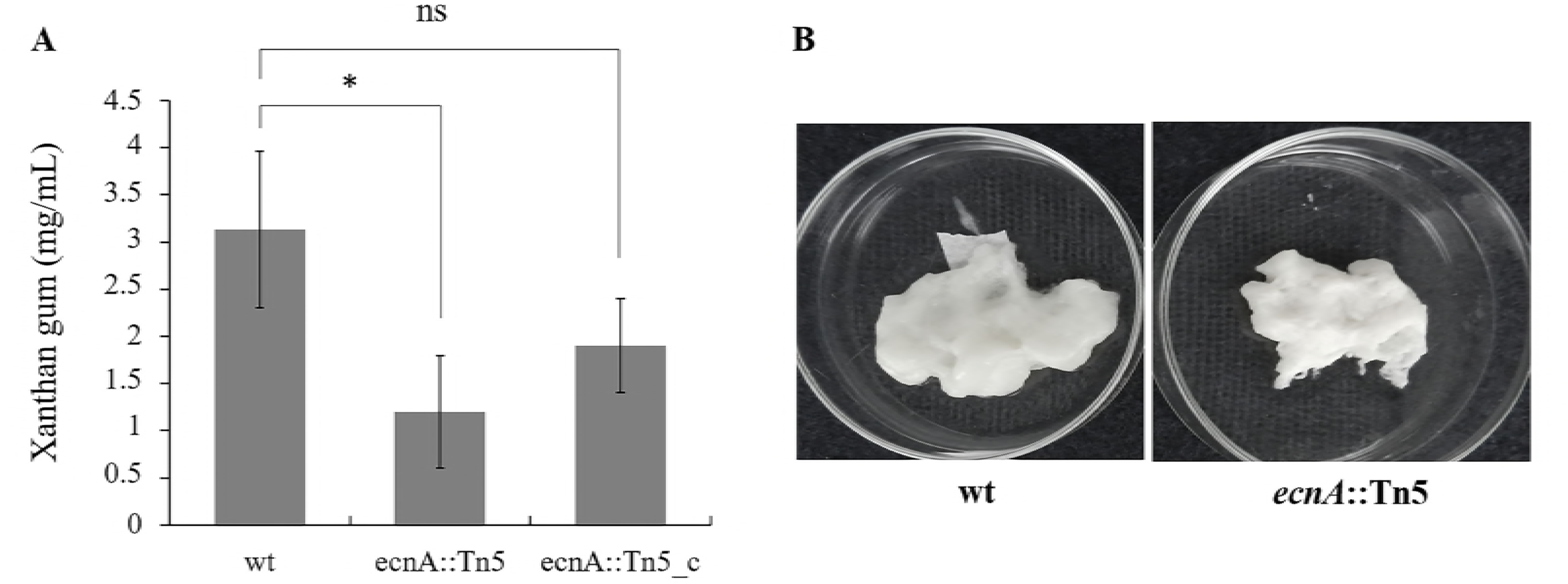
Effect of *ecnA* mutation in xanthan gum production. (A) Xanthan gum production by wild type, *ecnA* mutant and complemented strains. The data are the means of triplicate measurements from a representative experiment; similar results were obtained in two other independent experiments. *Indicates significant difference (*P* < 0.05) and ns indicates no significance. Error bars indicate the standard deviation of the mean. (B) Crude xanthan gum from wild-type and the dryer xanthan aspect from *ecnA* mutant. wt = wild-type *X. citri, ecnA*::Tn5 = *ecnA* mutant, *ecnA*::Tn5_c = complemented *ecnA* mutant.

**Figure 6.**
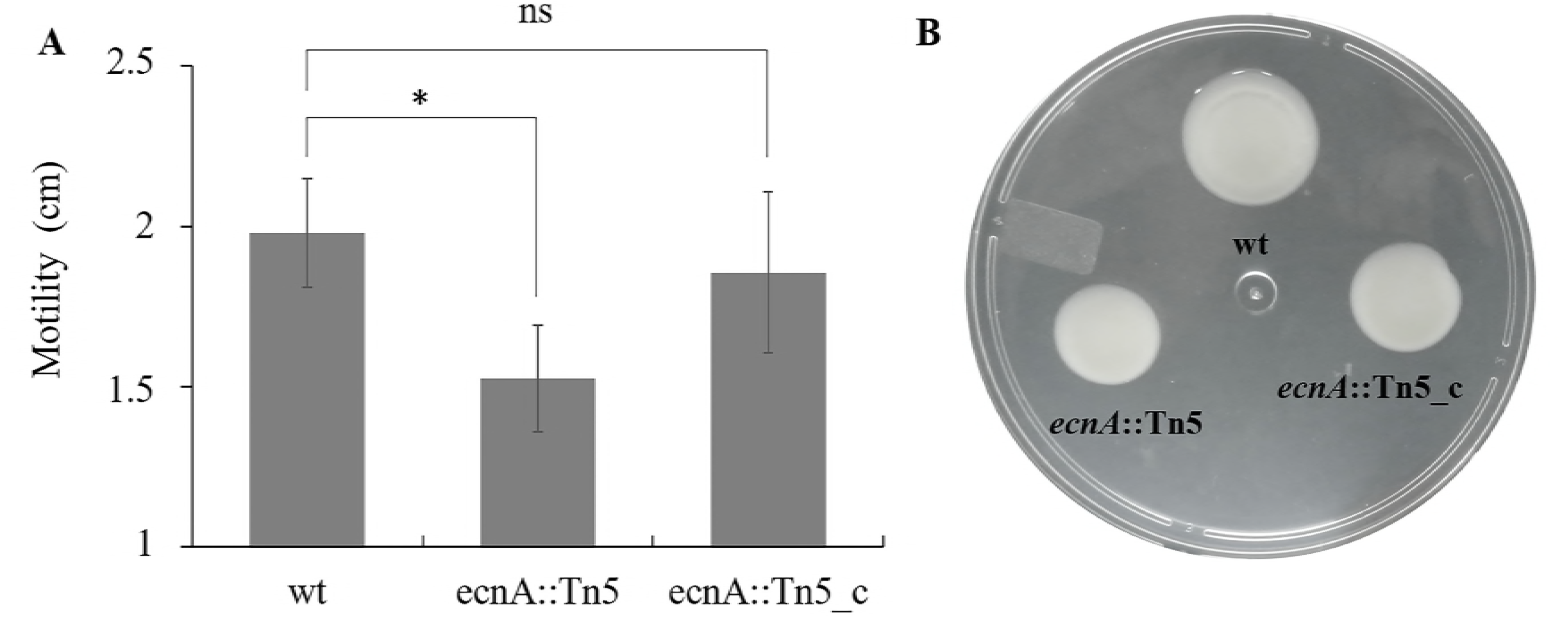
Effect of *ecnA* mutation in sliding motility. (A) Diameters of cell sliding on plates. *Indicates significant difference (*P* < 0.05) and ns indicates no significant. Error bars indicate the standard deviation of the mean. (B) *ecnA*::Tn5 showed an increase in motility that could be restored to wild-type levels by the introduction of a plasmid containing the intact *ecnA* gene. Cells from bacterial cultures at exponential stage were spotted on SB plates supplemented with 0.5% agar. Plates were incubated at 28 °C and photographed after 2 days. wt = wild-type *X. citri* subsp. *citri* strain 306, *ecnA*::Tn5 = *ecnA* mutant, *ecnA*::Tn5_c = complemented *ecnA* mutant.

### *ecnA* is involved in virulence and bacterial survival in plant host

The pathogenicity of the *ecnA* mutant was evaluated by infiltration and spray on the plant host. The bacterial population of the *ecnA* mutant in infiltrated leaves was significantly lower than wild-type at 21 dai (days after inoculation) (Fig. 7A), showing 5 logs difference. In addition, the symptoms were clearly reduced compared to the leaves infiltrated with wild-type (Fig. 7B).

**Figure 7.**
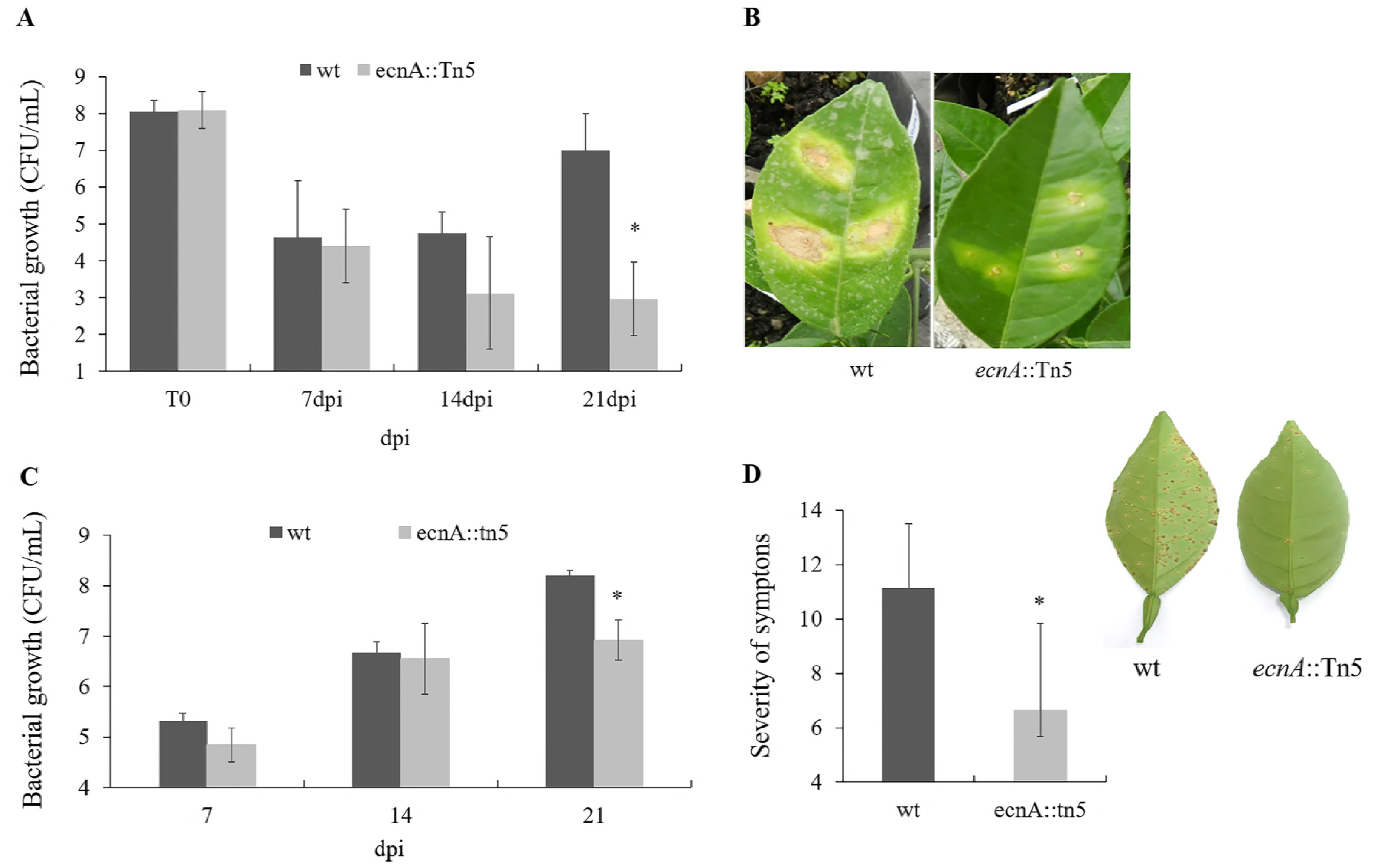
Bacterial growth and pathogenicity assay in planta. (A) Bacterial populations of wild-type and *ecnA* mutant inoculated by infiltration at a concentration of 10^8^ CFU/mL and analysed at 7, 14 and 21 days after inoculation (dai). (B) Symptoms of sweet orange leaves at 21 dai infiltrated with wild-type and *ecnA* mutant strains. (C) Bacterial populations of wild-type and *ecnA* mutant inoculated by spraying at a concentration of 10^8^ CFU/mL and analysed at 7, 14 and 21 dai. (D) Severity of citrus canker symptoms at 21 dai. Error bars indicate the standard deviation of the mean of two independent experiments. *Indicates significant difference *(P <* 0.05). Images are representative of five independent replicates at 21 dai.

When both bacteria were inoculated by spray, a method that better mimics the natural transmission condition, the *ecnA* mutant population was also significantly lower than that of the wild-type at 21 dai (Fig. 7C) resulting in a significant difference in symptoms severity of leaves (Fig. 7D). As *ecnA* is part of a TA system and it may be associated to tolerance to stress conditions (24, 46), we speculated that the lower bacterial population of *ecnA* mutant observed in plant could be due to a reduced tolerance to the plant defense response, such as reactive oxygen species (ROS) production.

### *ecnA* is involved with bacterial survival during oxidative stress

Bacterial populations were analyzed under ROS stress induced by hydrogen peroxide. The number of wild-type and *ecnA*+ cells significantly reduced (approximately 2 logs) after treatment (Fig. 8A), but the effect was much stronger for the *ecnA* mutant where no cultivable cells were recovered after stress. Besides, the relative expression of *ecnA* and *ecnB* was significantly repressed at high concentration of hydrogen peroxide (50mM) (Fig. 8B). These results are similar to previously published data on toxin-antitoxin in *X. citri*, where it was found that most stressor agents indeed repress the toxin-antitoxin systems in these bacteria (29). Altogether these results suggest that the lower population of *ecnA* mutant observed in the leaves could be a consequence of the less ability to cope with plant defense responses.

**Figure 8.**
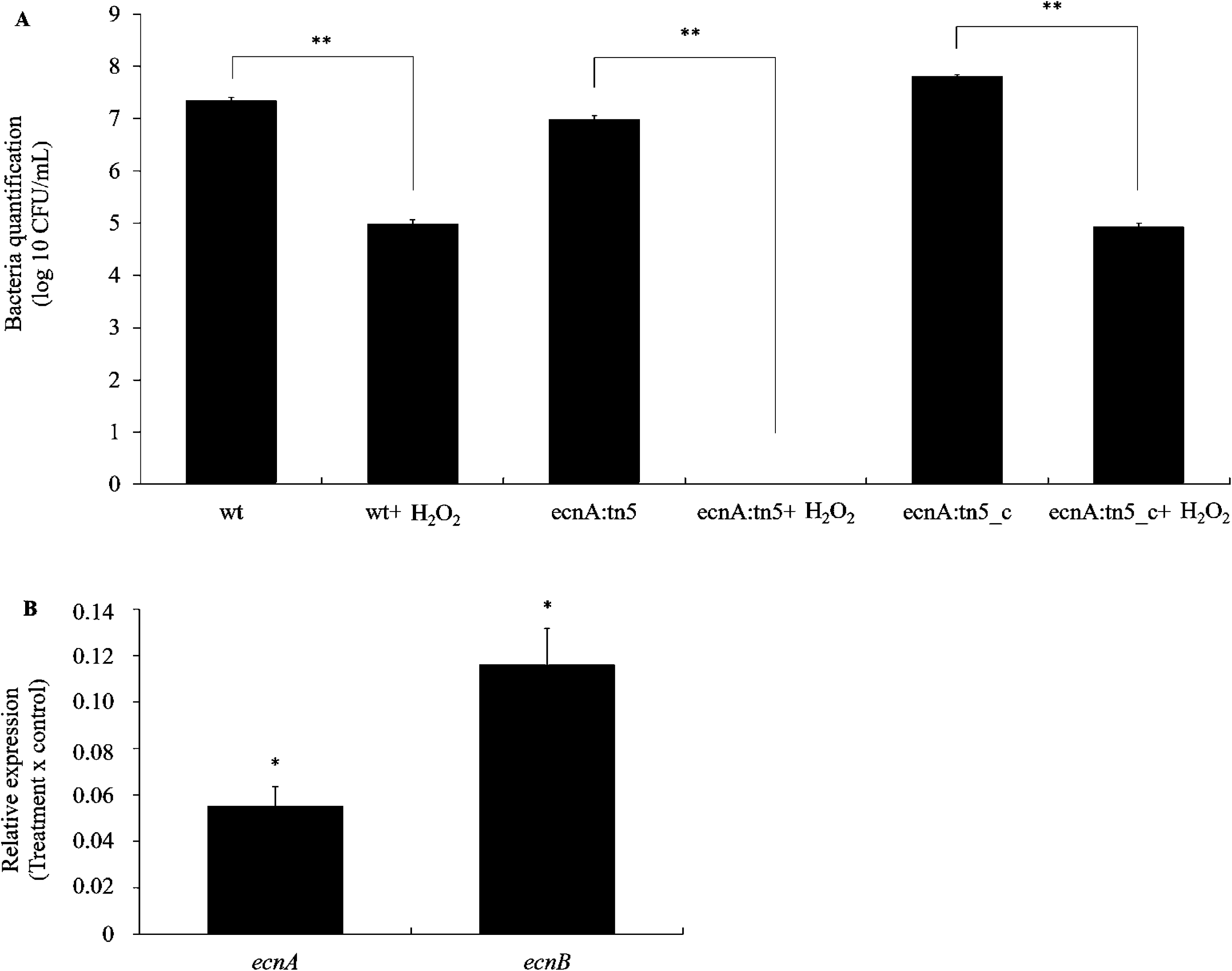
Sensitivity of wild-type and mutant strains to hydrogen peroxide (H_2_O_2_) **A.** Bacteria quantification (CFU/mL) before and after H_2_O_2_ stress. **B.** Relative expression of *ecnA* and *ecnB* in *X. citri* after H_2_O_2_ stress compared to cells without stress. The transcript abundance was determined by RT-qPCR. Data are shown as the mean of two independent biological experiments and three technical replicates. Error bars indicate the standard deviation of the mean. 16S rRNA was used as endogenous control. *Indicates significant difference *P* < 0.05 and ** *P* < 0.01.

## DISCUSSION

The *ecnAB* locus encodes two small peptides, named EcnA and EcnB (Fig. 1), and which has been considered a toxin-antitoxin system in *E. coli* because of similarities to the plasmid addiction module, *mazEF* (24). Sequences like *ecnAB* have been annotated in the *Xanthomonas* genomes (Fig. 2), indicating that this operon may play a conserved role. In *E. coli*, different functions have been proposed for the *ecnAB* TA system, including programmed cell death (24) and adherence in the respiratory tract of humans (26). In *Xanthomonas* it has been demonstrated that *ecnAB* is regulated by quorum sensing (18). Indeed our data show that *ecnA* and *ecnB* were down-regulated in the *rpfF* deletion mutant (Fig. 3), corroborating the transcriptome assay performed for regulatory analysis of the quorum-sensing system in *X. citri* (18). Curiously the biofilm formation phenotype is contrasting in the *ecnA* and *rpfF* mutants, while the *ecnA* mutation leads to reduced biofilm, the *rpfF* mutant increases the biofilm. This opposing phenotype is also observed for *gumB* since the mutant also presents less biofilm even though it is positively regulated by *rpfF* (45). These results could be explained by a negative regulation of adhesion proteins by *rpfF* in *Xanthomonas* (18) and therefore mutation in this gene would lead not only to a decrease in expression of *gumB* and *ecnA* but increase of genes related to adhesion. These later genes are responsible for the increased biofilm, which may not be under the regulation of *ecnA* or *gumB*, and that would explain the contrasting phenotype observed in these genetic backgrounds.

Our results suggest that the correlation of *ecnAB* and biofilm formation occurs due the EPS production as the *ecnA* mutant presents less biofilm and EPS (Fig. 4 and 5). Additionally, the *ecnA* mutant showed decreased sliding motility on the media surface (Fig. 6), which could affect the bacterial epiphytic growth and consequently the pathogenicity. It has been demonstrated that inefficiency of epiphytic growth leads to a less virulent behavior of *X. citri* (11, 40). This could explain the difference observed in symptoms development when the bacteria were sprayed but not infiltrated (Fig. 7). Curiously fewer symptoms were also observed after infiltration of the bacteria, which suggests that the role of *ecnAB* is not only associated with better fitness of epiphytical behavior but could also be involved in tolerance to plant defense responses. Bursts of ROS are quickly produced as a defense response to pathogen attack (47). Indeed, the *ecnA* mutant presents higher susceptibility for H_2_O_2_ (Fig. 8A), which may explain the less symptoms observed when the mutant is infiltrated together with reduction in cell population.

We show that *ecnA* and *ecnB* genes were down-regulated under hydrogen peroxide stresses (Fig. 8B). Similarly, the TA systems XACa0028/XACa0027 and *chpBS* were also significantly down-regulated by copper and temperature stresses in *X. citri* (29). The transcriptional repression of TAs under stress conditions suggests that the amount of unstable antitoxin in the cell reduces, and then the stable toxin is free and active (48, 49). The reasons for such repression are still elusive, but there is a possibility that it is involved in the persister cell formation, a phenotype recently explored for phytopathogens and its implication in environmental stress survival (50).

## ACKNOWLEDGEMENTS

This work was supported by a research grant from INCT Citrus (Proc. CNPQ 465440/2014-2 and FAPESP 2014/50880-0) and the Fundação de Amparo à Pesquisa do Estado de São Paulo (FAPESP - 2013/10957-0). L. M. Granato, a PhD. student from the Program in Genetics and Molecular Biology at the Institute of Biology at the State University of Campinas, was supported by a fellowship from CAPES/PSDE fellowship (99999.002657/2014–07). P. M. M. Martins is a post-doctoral fellows (FAPESP 2016/01273-9). M. A. Takita, M.M. Machado and A. A. De Souza are recipients of research fellowships from CNPq. The authors declare no competing or financial interests.

---

**Supplementary Fig 1.** Swimming motility *X. citri*, its *ecnA* mutant, and the complemented *ecnA* mutant. Cells were inoculated onto NYGB medium supplemented with 0.25 % agar and photographed after 2 days of incubation at 28 °C. wt = wild-type *X. citri* subsp. *citri* strain 306, *ecnA*::Tn5 = *ecnA* mutant, *ecnA*::Tn5_c = complemented *ecnA* mutant.

**Supplementary Fig 2.** Growth curve of *X. citri*, its *ecnA* mutant, and the complemented *ecnA* mutant on NBY medium. Optical density (OD) at 600 nm was analysed every 2 h by Varioskan Flash (Thermo Scientific). There is no significant difference between the strains. wt = wild-type *X. citri* subsp. *citri* strain 306, *ecnA*::Tn5 = *ecnA* mutant, *ecnA*::Tn5_c = complemented *ecnA* mutant.

